# Fatty Alcohols, a Minor Component of the Tree Tobacco Surface Wax, Reduce Insect Herbivory

**DOI:** 10.1101/2021.07.15.452450

**Authors:** Boaz Negin, Lior Shachar, Sagit Meir, Claudio C. Ramirez, A. Rami Horowitz, Georg Jander, Asaph Aharoni

**Affiliations:** Plant and Environmental Science Department, Weizmann Institute of Science, Rehovot 7610001, Israel; Centre for Molecular and Functional Ecology in Agroecosystems, Instituto de Ciencias Biológicas. Universidad de Talca, 2 Norte 685, Talca, Maule, Chile; Department of Entomology, Agricultural Research Organization (ARO), Gilat Research Center for Arid & Semi-Arid Agricultural Research, 85280, Israel; and Katif Research Center, Sedot Negev; Ministry of Science and Technology 85200, Netivot, Israel; Boyce Thompson Institute, Ithaca, NY 14853, USA

## Abstract

Despite decades of research resulting in a comprehensive understanding of epicuticular wax biosynthesis and metabolism, the function of these almost ubiquitous metabolites in plant-herbivore interactions remains unresolved. To develop a better understanding of this role, we investigated plant-herbivore interactions in four *Nicotiana glauca* (tree tobacco) genome edited mutants. This included [*eceriferum1* (*cer1)*, *eceriferum3* (*cer3), β-ketoacyl-coA synthase6* (*kcs6*), and *fatty acyl-coA reductase* (*far*)] displaying a wide range of alkane and fatty alcohol abundances. Three interaction classes were examined: chewing herbivory with seven caterpillar and one snail species, phloem feeding with *Myzus persicae* (green peach aphid), and egg laying with *Bemisia tabaci* (sweet potato whitefly). We found that high wax load and alkane abundance did not reduce caterpillar or snail herbivory. However, fatty alcohol content was negatively correlated with caterpillar growth, suggesting a role in reducing insect herbivory despite its lower levels. Aphid reproduction and feeding activity were not correlated with wax load and composition but are potentially affected by altered cutin composition of *cer1* mutants. When examining non-feeding activities, wax crystal morphology could explain the preference of *B. tabaci* to lay eggs on wildtype plants relative to *cer1* and *far* mutants. Accordingly, the fatty alcohol wax component reduces caterpillar herbivory on the chemical level, but oviposition is increased when wax crystals are dense. The results suggest that this varied response between herbivore classes and species, at times displaying increased and at times reduced fitness in response to altered wax composition is in part a consequence of co-evolution that shaped the specific effects of different *N. glauca* metabolites such as anabasine and fatty alcohols in plant-herbivore interactions.

## Introduction

An epicuticular wax layer coating of above-ground organs is distributed almost ubiquitously in the plant kingdom (Kong et al., 2020), including in mosses (Proctor, 1979), ferns (Guo et al., 2018), gymnosperms (Oros et al., 1999), and angiosperms (Lee and Suh, 2015). Decades of research on plant epicuticular waxes have led to a comprehensive understanding of their biosynthesis, transport, and regulation (Aharoni et al., 2004; Seo et al., 2011; Daszkowska-Golec, 2020). The epicuticular wax layer is part of a complex structure of the plant cuticle together with the cutin polymer, which is embedded with both intracuticular waxes and phenolic compounds asides from its fatty acid backbone. The cutin matrix is formed from C_16_-C_18_ fatty acids (FA) that can be hydroxylated either at the FA’s end (ω-hydroxylated), throughout the remaining carbons, contain a dual carboxylic group or create epoxy bonds. These cutin monomers are then esterified at their functional groups creating an amorphic matrix, incorporating glycerol, phenols and wax components [reviewed in Pollard et al., (2008); Schreiber, (2010)]. Given both its structure and function, cutin metabolism is strongly connected to wax metabolism through common precursors, transcriptional regulation (Oshima et al., 2013) and biosynthetic enzymes (Lü et al., 2009).

Production of epicuticular waxes begins with fatty acid synthesis in plastids (Fig. 1). These are then transferred to the endoplasmic reticulum (ER) where the C_16_ and C_18_ FA are converted to acyl-CoA by LONG CHAIN ACYL SYNTHETASE (LACS; Schnurr et al., 2004; Lü et al., 2009) and elongated by the fatty acid elongase complex. This complex includes four enzymes: β-ketoacyl-CoA synthase (KCS; Millar and Kunst, 1997), which has different variants responsible for elongation to different carbon chain lengths (Hegebarth and Jetter, 2017). The *KCS6* gene examined in this study, for example, is responsible for chain elongation from 24 to 34 carbons (Millar et al., 1999). β-Ketoacyl-CoA reductase (KCR; Beaudoin et al., 2009), 3-hydroxyacyl-CoA dehydratase (HCD; Bach et al., 2008), and trans-2,3-enoyl-CoA reductase (ECR; Zheng et al., 2005) are other components of an elongation cycle that produces a mix of very long chain fatty acyl-CoA with different chain lengths. Acyl-CoAs next enter the alkane or the primary alcohol forming pathways. In the alkane pathway, acyl-CoAs are decarboxylated to form aldehydes and then alkanes through a yet unidentified mechanism facilitated by the CER1 and CER3 enzymes (Aarts et al., 1995; Chen et al., 2003; Bourdenx et al., 2011). Alkanes may be further modified to form secondary alcohols and ketones by a cytochrome p450 MID-CHAIN ALKANE HYDROXYLASE1 (MAH1), which catalyzes both reactions (Greer et al., 2007). In the primary alcohol pathway, very long chain acyl-CoAs are converted to primary alcohols by addition of a hydroxyl by the FATTY ACYL-COA REDUCTASE (FAR) enzyme (Rowland et al., 2006). An additional stage in this pathway is the conjugation of C_16_ or C_18_ fatty acids to the primary alcohols to form wax esters by the bifunctional wax synthase/ acyl-CoA:diacylglycerol acyltransferase1 (WSD1; Li et al., 2008).

**Figure 1.**
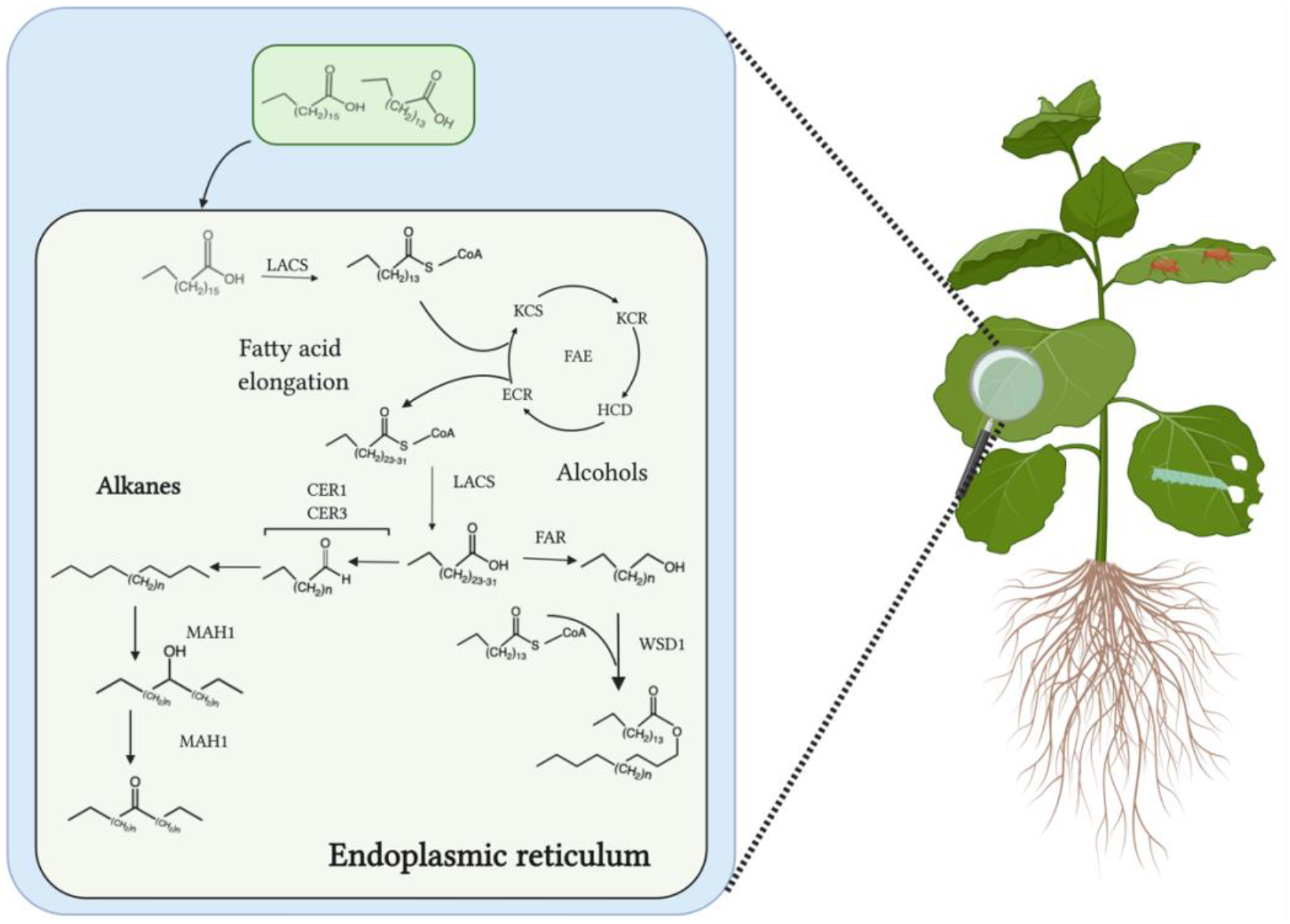
Schematic representation of the major epicuticular wax synthesis pathways. Here we characterized mutants generated in a previous study (Negin et al., 2021) that were genome edited, including the *CER1*, *CER3*, *KCS6* and *FAR* genes.

Following their synthesis, wax monomers are transported from the ER and secreted beyond the cutin layer where they form crystal-like structures. These crystals are shaped by their monomer composition and similar wax synthesis mutants from different species display similar morphological changes in their wax crystal structure (Koornneef et al., 1989; Negin et al., 2021). This conservation hints to the crystals being self-assembled, but whether an enzymatic process is involved non-the less remains to be determined (Koch and Ensikat, 2008).

To date, the specific fitness advantages of plant epicuticular waxes have not been fully investigated. Among the many functions that have been suggested (Barthlott and Neinhuis, 1997; Long et al., 2003; Bourdenx et al., 2011), the most prominent is the prevention of water loss (Clarke and Richards, 1988; Islam et al., 2009). However, it is not known whether epicuticular wax is the primary factor contributing to desiccation resistance or whether other cuticular layers such as the cutin proper or intracuticular wax are more important (Riederer and Schreiber, 2001). Additionally, as the first barrier that most herbivores encounter, epicuticular wax has been suggested to play a role in plant-insect interactions. The specific function is dependent on the feeding habits of the attacking insects: chewing (Johnson and Severson, 1984; Saleem et al., 2019; Bernaola et al., 2021), boring, or phloem feeding (Bergman et al., 1991; Žnidarčič et al., 2008; Wójcicka, 2013). Additionally, oviposition (Udayagiri and Mason, 1997; Li and Ishikawa, 2006; Rid et al., 2018) or the construction of living spaces may be affected by leaf surface waxes. A compelling example of the complex interaction between epicuticular wax and insect interactions can be found in macaranga trees. In these, stems are covered by a slippery wax layer that selectively allows the climbing only of a symbiotic ant species (Federle et al., 1997; Markstädter et al., 2000). However, in general, a glossy phenotype with reduced epicuticular waxes is correlated with greater resistance to insect attack (Eigenbrode and Espelie, 1995; Shepherd et al., 1999), and even when reducing the adherence of insects to plant surfaces and thus disrupting foraging, slippery surfaces may aid herbivores by reducing their susceptibility to predation (Eigenbrode, 2004). Furthermore, epicuticular wax components have been found to stimulate feeding in some insect species (Mori, 1982; Karmakar et al., 2016).

*Nicotiana glauca* is a perennial shrub native to south America, but which has since spread worldwide. Stems and leaves of *N. glauca* have a grayish glaucous appearance, a striking phenotype that gave this species its name. The glaucous appearance is caused by a dense epicuticular wax crystal layer, which has a relatively simple composition dominated by a C_31_ alkane with lesser amounts of primary alcohols and additional alkanes. The combination of a high wax load, simple wax composition, a high quality genome sequence (Usadel et al., 2018), and an established transformation system that can be used for CRISPR/Cas9 mutagenesis, make *N. glauca* a perfect model system for epicuticular wax research. In addition, the ability of *N. glauca* to grow under diverse conditions, including growth rooms, greenhouses, and natural environments worldwide, makes it fitting for the study of interactions with insect herbivores.

To investigate the effects of plant surface waxes on insect herbivory, we utilized a recently generated set of *N. glauca* mutants that had cuticular lipids metabolism genes knocked out using CRISPR/Cas9 [described at length in Negin et al. (2021)]. Specifically, we conducted experiments with four mutants carrying mutations in epicuticular wax biosynthesis genes: *cer1*, *cer3, kcs6*, and *far*. Caterpillar and snail feeding, *Myzus persicae* (green peach aphid) reproduction and feeding patterns, and *Bemisia tabaci* (sweetpotato whitefly) egg laying were examined in wild type and epicuticular wax-deficient *N. glauca* plants. We discovered that leaf chewing was not affected by alkane abundance, but rather by the far less abundant fatty alcohol component. Phloem feeders were not affected by wax composition, but rather by cutin disruption while oviposition was affected by wax crystal structure but not chemical composition.

## Materials and methods

### Plant material and growth conditions

*Nicotiana glauca* seeds were collected from plants growing near the Weizmann Institute campus. Plants used for epicuticular wax extraction were grown in a growth room at 22°C with a 16:8 h light:dark cycle, in potting soil (Even Ari; http://www.evenari.co.il/), consisting of peat, coconut fiber, quartz and slow release fertilizer. Plants used for assessment of anabasine and sugar content were grown in a greenhouse during the winter and spring seasons with ambient lighting in soil containing peat, tuff and slow-release fertilizer. Plants used for caterpillar, aphid, and whitefly egg laying assays were grown in rooms as for wax extraction. Leaves used for the snail herbivory assays were grown in a greenhouse during the winter months.

### Snails and Insects used in the Experiments

*Helix engaddensis* snails were collected from areas next to the tree tobacco plots near the Weizmann institute campus. *Spodoptera exigua* (fall armyworm), *Spodoptera eridania* (southern armyworm), *Helicoverpa zea* (corn earworm), *Trichoplusia ni* (cabbage looper), and *Heliothis virescens* (tobacco budworm) eggs were obtained from Benson Research (https://www.benzonresearch.com/), hatched on beet armyworm diet (www.southlandproducts.net), and placed on plant leaves as first-instar larvae after 2-3 days. *Manduca sexta* (tobacco hornworm) eggs were kindly supplied by Robert Raguso (Department of Neurobiology and Behavior, Cornell University). *Spodoptera littoralis* (Egyptian cotton leafworm) larvae and *M. persicae* (green peach aphids) were provided by Metabolic Insights Ltd (Ness Ziona, Israel). The latter were raised on either pepper or tree tobacco plants in a growth room at 20°C under continuous lighting. A tobacco-adapted red strain of *M. persicae*, which has been described as the “USDA” strain (Ramsey et al., 2007; Ramsey et al., 2014), was maintained on *Nicotiana tabacum* (tobacco) plants in a growth room at 23°C with a 16:8 h light:dark cycle. *B. tabaci* (whiteflies), which were originally collected from cotton fields during the 2010 season, have been reared on cotton seedlings, (*Gossypium hirsutum L)* under standard conditions of 26±1°C, 50% R.H., and a 14:10 h light:dark cycle. Since being collected, this population has been isolated and not treated with any insecticides. This colony belongs to B biotype (or MEAM1 species) of *B. tabaci.*

### Wax composition profiling

Epicuticular wax was extracted from plants raised in a growth room. Three 12 mm diameter discs were taken from leaves and dipped in 4 ml chloroform with 10 μg of a C36 alkane internal standard, shaken gently for 15 seconds, the chloroform extract was transferred to 1 ml reaction vials (Sigma Aldrich, https://www.sigmaaldrich.com/) and evaporated under a nitrogen flow. Samples were resuspended in 200 μl chloroform to which 20 μl pyridine and 20 μl N,O-bis(trimethylsilyl)trifluoroacetamide (BSTFA) were added in each vial. Samples were derivatized at 70°C, and 3 μl were injected in a splitless mode into an Agilent Gas chromatography mass spectrometry (GC-MS) system (Agilent 7890A chromatograph, 5975C mass spectrometer and a 7683 auto sampler; Agilent, Santa Clara, U.S.) as described previously (Cohen et al., 2019). Results were analyzed using the Agilent ChemStation software, and metabolites were identified by comparison to fragmentation data in the NIST mass spectral library (https://chemdata.nist.gov/), as well as in-house standards and retention indexes. Quantification was performed using the Agilent ChemStation software by quantifying the peak area of a target ion, which was then normalized to the internal standard peak area.

### Polar compound profiling

Polar compound profiling was performed using GC-MS. Leaf discs with a 12 mm diameter were cut from three leaves of *N. glauca* plants and frozen in liquid nitrogen. Samples were ground to a fine powder and 100 mg were used for sample preparation. Extraction, derivatization, and GC-MS were performed as described previously (Korenblum et al., 2020), with 12 μg ribitol added to each of the samples as an internal standard. Data analysis, including metabolite identification and quantification, were performed using the Xcalibur software (Thermo Fisher Scientific; https://www.thermofisher.com/). Metabolites were identified by comparing ion fragmentation to the NIST mass spectral library and in-house standards and retention indexes. Once peaks were quantified, peak area was normalized to the ribitol peak area of the same sample.

### Caterpillar growth and snail feeding experiments

In one assay, caterpillars were confined to single leaves of six-week-old wild type (WT) *N. glauca*, or to leaves of *cer1*, *cer3*, *kcs6*, or *far* mutants in a growth room. Three leaves from each plant were individually inserted to organza mesh bags (www.amazon.com, item B073J4RS9C). A single neonate was placed on each leaf and left to feed for 9-11 days (*T. ni* and *M. sexta* fed for 9 days, *S. exigua*, *H. virescens* and *H. zea* fed for 10 days, and *S. eridania* fed for 11 days). Following this, caterpillars were removed from the bags, and survival rates and weight were recorded. For experiments with caterpillars on whole plants, WT *N. glauca*, and wax mutants were raised in a growth room for six weeks. Ten *S. littoralis* neonates were placed in mesh cages containing two wildtype plants or two wax mutant plants from the same genotype. The caterpillars were allowed to feed on the plants for 12 days, after which they were weighed. *Helix engaddensis* snails were maintained on lettuce for several days, and starved for 24 h before the experiment. Snails were placed to 10.5 cm x 10.5 cm x 9.5 cm Magenta boxes that contained a wildtype *N. glauca* leaf and either a *cer3* mutant leaf or a *kcs6* mutant leaf. Leaves placed in each Magenta box were of a similar size, which ranged between 40-60 cm^2^. The leaves were photographed prior to being placed in the Magenta boxes and again following 48 h of snail feeding. The leaf areas were analyzed using ImageJ software (https://imagej.nih.gov/ij/) and the leaf area consumed was calculated. This assay was repeated three times in a similar manner.

### Aphid reproduction and monitoring of aphid feeding activity

Wild type *N. glauca*, and wax mutants were raised in a growth room for six weeks, after which three *M. persicae* aphids were caged on the adaxial side of individual leaves. These plants were placed in a growth room at 22°C with a 16:8 h light:dark period. Aphids were then counted approximately every two days for a duration of 14 days. This assay was repeated three times with aphids that were previously raised on *N. glauca* and five times with aphids that had been raised on *Capsicum annuum* (pepper) plants. Aphid feeding behavior was monitored using an electric penetration graph (EPG) system (https://www.epgsystems.eu/; Tjallingii and Esch, 1993). Apterous adult asexual females of *M. persicae* were connected to a 25-μm-thin gold wire using a droplet of conductive silver glue (PELCO Colloidal Silver No 16031; Ted Pella, Red- ding, CA, USA) on their abdomen. Aphids were starved for one hour prior to being connected to the EPG system. Once all aphids in a replicate were attached to the system, they were simultaneously lowered on to leaves of either WT or *cer1* plants, where they were left for eight hours, and their activity was recorded continuously. The system recorded the following waveforms, which were correlated with known activities: NP – non probing phase, when the aphid stylet is not inserted inside the leaf; C – activities during penetration which are mainly intercellular (probing); E1 – salivation in a sieve element, E2 – ingestion of fluids from the phloem; F – derailed stylet mechanisms; G – xylem derived ingestion, and PD – potential drops which indicate the stylet moved from being positioned intercellularly to being positioned within a cell. The EPG waveforms were recorded using Stylet+d software (EPG Systems) and analyzed using A2EPG software (Adasme-Carreño et al., 2015). The sequences of waveforms for each replicate were imported into the Excel workbook of Sarria *et al.* (2009) to automatically calculate all the sequential and non-sequential EPG variables that characterize probing and ingestion phases.

### Bemisia tabaci egg laying

Wildtype *N. glauca*, and wax mutants were raised in a growth room for six weeks, after which the plants were used for insect assays. Cages were attached to abaxial sides of individual leaves, with each cage on an individual plant. Twenty adult *B. tabaci* were placed in each cage for 48 h. Leaf cages were removed and the number of eggs laid was counted using a binocular microscope. This assay was repeated three times.

### Statistical analysis

Student’s *t-*tests were used for pairwise comparisons of mutant lines when compared to WT *N. glauca* for wax analysis, which was performed six times of which a representative dataset was selected. In other cases of multiple comparisons with a WT control, Dunnett’s test was used. The exception to this was that, when comparing data obtained using the EPG system, a non-parametric Wilcoxon test was used. When comparing different caterpillar species weight and survival percentage, Tukey’s HSD and Chi-square tests were used respectively. To determine whether epicuticular wax mutations had a significant effect, without relation to the affected gene, a one-way ANOVA was performed prior to the analyses of the individual line effects. A two-way ANOVA was used in the aphid reproduction assay to determine if there was a significantly different response of the various genotypes to the passage of time. These statistical analyses were performed using JMP 14 software (SAS Institute; http://www.jmp.com/en_us/home.html). A single asterisk represents significance of p < 0.05 and double asterisks represent a significance of p < 0.01 when compared to WT *N. glauca*. In the Tukey’s HSD and Chi square tests, different letters represent a significance of p < 0.05. In addition, F tests used to determine regression line slope equality to zero were performed using GraphPad software (https://www.graphpad.com/).

## Results

### Herbivory on N. glauca plants grown in natural settings

To study the impact of altered epicuticular wax composition on *N. glauca* plant herbivory we first examined the insects inhabiting this wild species in nature. *N. glauca* plants grown in natural settings were populated by various herbivores including snails (*H. engaddensis*), caterpillars (*Vanessa cardui* and *Ocnogyna loewii*), and leaf aphids. We observed several classes of biotic interactions. The most apparent was leaf chewing, which involved both caterpillars (Fig. 2A-B) and snails (Fig. 2C) that caused substantial damage to leaves. The second class included phloem-feeding insects, in the form of aphids that colonized, reproduced, and fed on the plants (Fig. 2D). The third and most varied class involved the use of *N. glauca* for non-feeding interactions such as egg-laying, attaching pupae, building nests for larvae, and other activities (Fig. 2E-F). These varied relations, monitored under natural settings, led us to examine the three interaction classes in more detail under controlled laboratory conditions.

**Figure 2.**
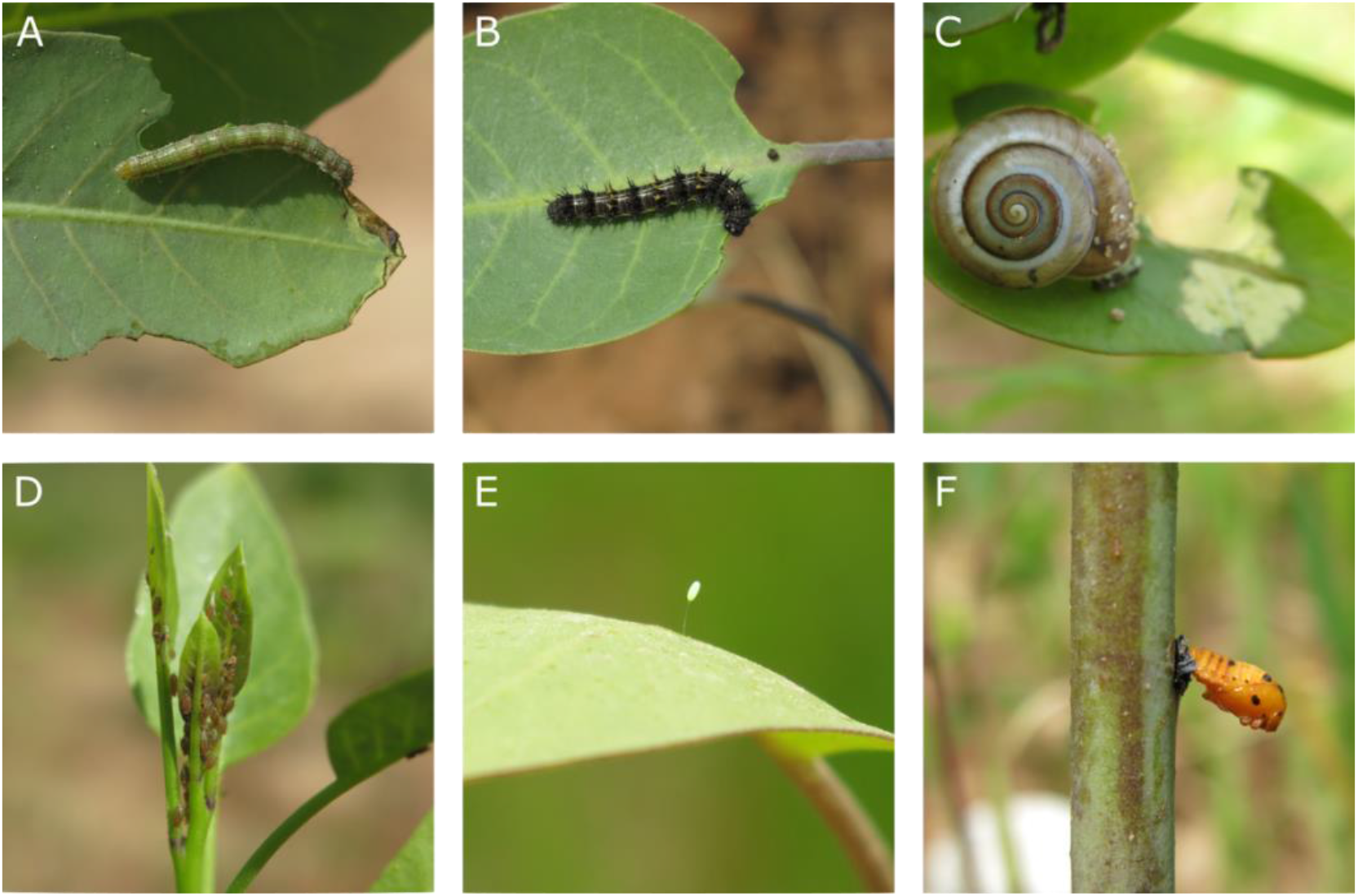
Herbivore – plant interactions observed in *N. glauca* grown in a natural setting. (**A-C)** Chewing herbivores. (**D)** Phloem feeders. (**E-F)** Non-feeding interactions. In (A), putatively identified *Spodoptera exigua* caterpillar feeding on a *N. glauca* leaf; (B) a *Venesa cardui* caterpillar feeding on a *N. glauca* leaf; (C) *H. engaddensis* on a *N. glauca* leaf after having fed on it; (D) leaf aphids clustering on the shoot apex; an insect egg, possibly green lacewing (*Chrysoperla rufilabris*) laid on a *N. glauca* leaf; (F) the pupa of *Coccinella septempunctata* attached to the stem of *N. glauca*.

### Exploiting a set of cuticular wax mutants to study plant-herbivore interactions

Very recently, we employed CRISPR-based genome editing in *N. glauca* and generated 16 mutants altered in cuticular lipids metabolism genes (Negin et al., 2021). In the plant-insect interaction assays in this study, we concentrated on four wax biosynthetic mutants contained within this set, including *eceriferum1 (cer1), eceriferum 3 (cer3)*, *β*-ketoacyl-CoA synthase 6 (*kcs6*) and *fatty acyl-coa reductase (far).* Our earlier work examined cuticular wax composition in *N. glauca* plants growing under greenhouse conditions, which may have had a different impact on wax metabolite composition as compared to plants raised in growth rooms. Thus, in this study we analyzed epicuticular wax composition in leaf extracts derived from plants growing in parallel to those used for plant-insect interaction assays carried out in growth rooms.

Wild type (WT) *N. glauca* leaf epicuticular wax composition is typically dominated by a C_31_ alkane with lesser but substantial amounts of C_33_ alkane and trace amounts of C_29_, C_30_, and C_32_ alkanes (Fig. 3A-B). WT surface wax also contains substantial amounts of C_24_, C_26_, and C_28_ primary alcohols and C_26_ aldehydes (Fig. 3C-F). In *cer1* mutants, affected in alkane biosynthesis, alkane abundance is significantly reduced (Fig. 3A-B), whereas primary alcohols and aldehydes are less affected. Moreover, these mutants exhibit a significant (p=0.0003) increase in very long chain fatty acid content. Although *cer3* mutants are also affected in their alkane biosynthesis, we found a much less drastic effect on wax composition in these lines; the C_31_ alkane load is reduced to approximately 18% of WT, while the far less abundant C_33_ alkane load is increased 2.5 times as compared to control plants (Fig. 3A-B). The *kcs6* plants mutated in one of the four fatty acid elongase complex genes (see Fig. 1), display a higher abundance of wax components with low chain lengths and a reduced abundance of longer wax components. We found dramatically reduced alkane and C_26_ alcohols load in these plants (Fig. 3A-B and Fig. E). We also noted a significant rise in wax ester abundance, which may stem from a higher abundance of shorter primary alcohols. The *far* mutants affected in the primary alcohol-forming pathway, showed a most significant reduction in aldehydes and primary alcohol levels (Fig. 3C-F). The same plants possess a consistent trend of increased alkane load (Fig. 3A). Similar alterations in surface wax composition greatly affected wax crystal morphology which differed between the four mutant lines in previous studies (Negin et al., 2021).

**Figure 3.**
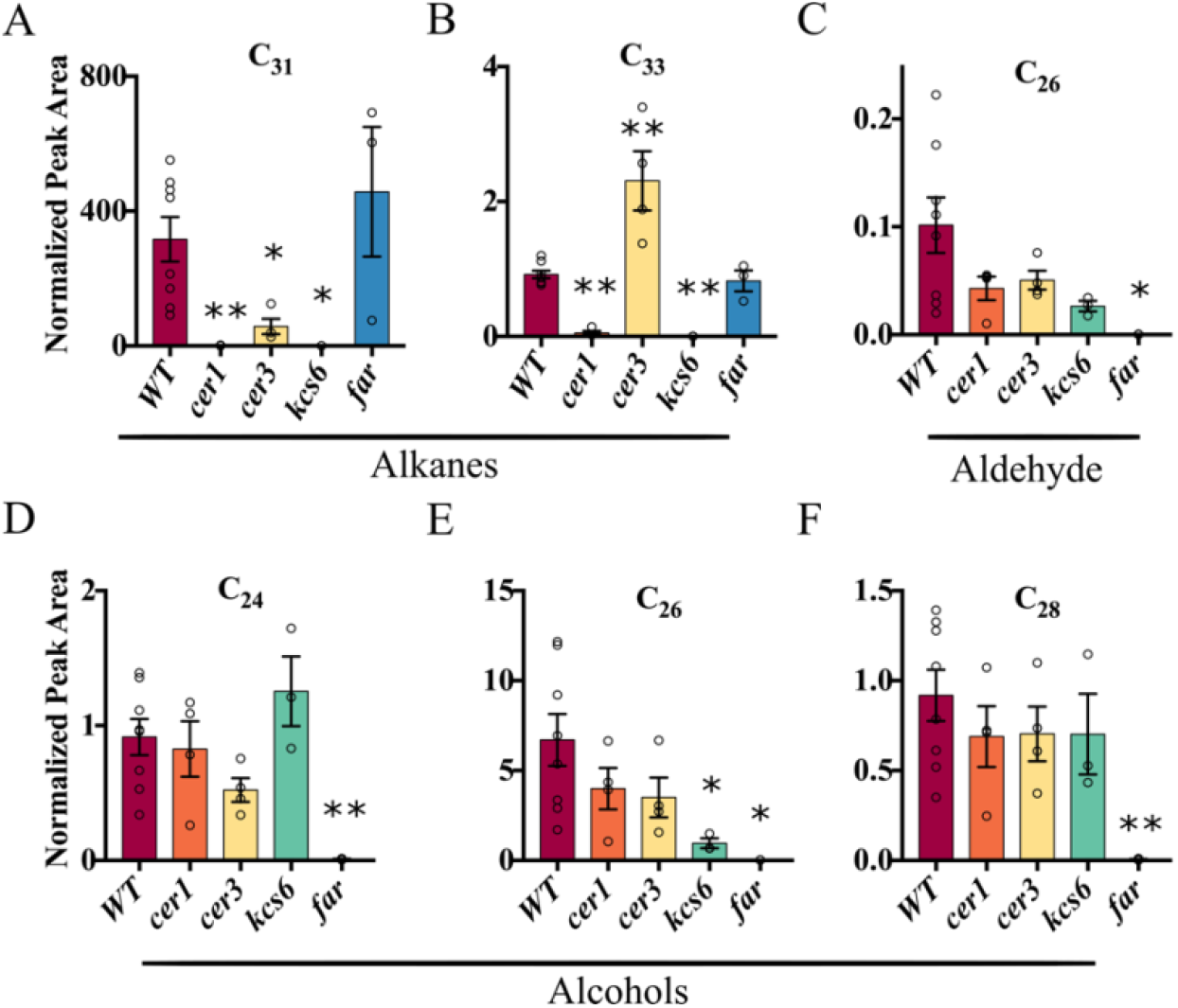
Relative abundance of the most prominent epicuticular wax components found in leaves of four growth room raised epicuticular wax metabolism mutant lines of *N. glauca* as compared to WT **(A)** C_31_ alkane. **(B)** C_33_ alkane. **(C)** C_26_ aldehyde. **(D)** C_24_ primary alcohol. **(E)** C_26_ primary alcohol. **(F)** C_28_ primary alcohol. Single asterisks indicate significance of p<0.05 while double asterisks indicate p<0.01 as determined in a Student’s *t*-test. WT, *cer1* and *cer3* n=4 *kcs6* and *far* n=3. WT = wildtype *N. glauca.*

In addition to wax composition, we also used Gas Chromatography Mass Spectrometry (GC-MS) and derivatized extracts to analyze the levels of polar metabolites in the different genotypes under study. These included anabasine and several highly abundant sugars. Anabasine is an alkaloid that is a structural isomer of nicotine and is known to be similarly toxic to insects (Zammit et al., 2014). Thus, we wanted to account for the effects of mutations in wax-related genes on the abundance of this metabolite in the mutants. While anabasine content varied between the genotypes, as determined by one-way ANOVA (P < 0.05), this was not converted to significance in any of the individual mutants examined (Fig. S1A). Sugar content varied widely as well, though the most abundant sugars did not reach significance (Fig. S1B-F).

### The impact of altered wax composition on chewing herbivores

To assess the effects of wax composition on chewing herbivores, we initially examined the feeding responses of six caterpillar species. Since little is known regarding caterpillar feeding habits on *N. glauca*, five broad generalists (*S. exigua*, *S. eridania*, *T. ni*, *H. zea*, and *H. virescens*) and one tobacco specialist (*M. sexta*) were used in these experiments. We found large variance in the average weight of the caterpillars, ranging from 16.4 mg in *H. zea* to 122.4 mg in the tobacco specialist *M. sexta* (Fig. 4A). Caterpillar survival rates varied widely as well. Only 38.5% of *H. zea* were alive upon weighing, whereas 91.5% of *T. ni* were alive at that point (Fig. 4B). Given these results, *H. zea* was left out of further analysis, to avoid confounding effects stemming from its incompatibility with the host plant. ANOVA analysis of the influence of mutations in wax genes on caterpillar weight revealed a significant effect (Fig. 4C). Furthermore, caterpillars feeding on *far* and *cer3* mutants were significantly heavier than those on *kcs6* and *cer1* mutants, although all mutant lines were statistically similar to WT due to its intermediate values (Fig. 4C). When we analyzed individual caterpillar species, those grown on the *far* and *cer3* mutants stood out; caterpillars feeding on them displayed a trend of elevated weight in three species (Fig. 4D, F-G). Moreover, caterpillars that fed either on *cer3* (Fig. 4E) or *far* (Fig. 4H) leaves were heaviest in the remaining species, though this was not significant when compared to WT plants.

**Figure 4.**
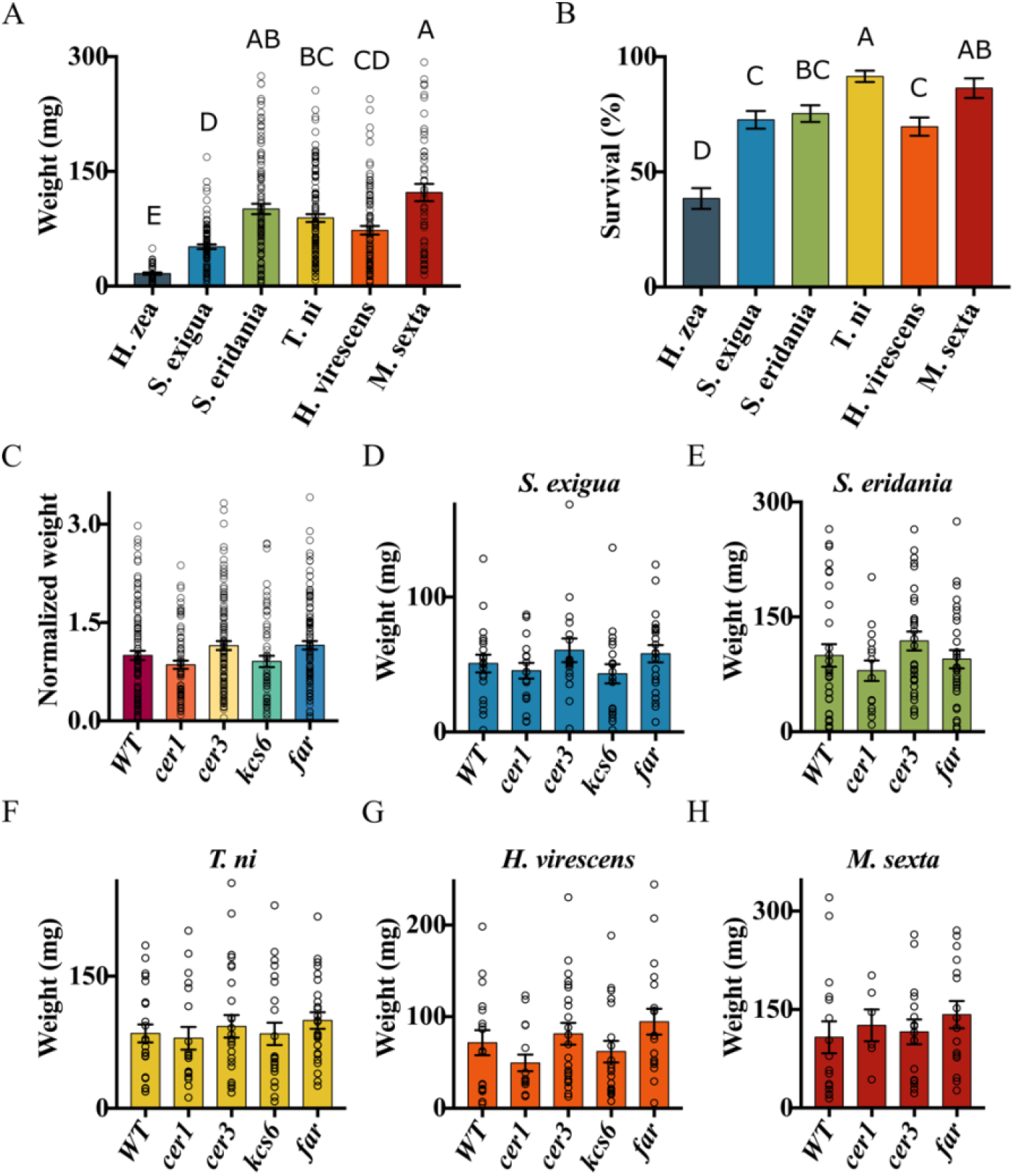
Six caterpillar species survival and growth on *N. glauca* leaves, and the effect of wax composition on growth of five of those species. Caterpillar weight **(A)** and survival rate **(B)** following 9-11 days of feeding on *N. glauca* leaves. Data combines results of both caterpillars grown on WT leaves and those grown on wax mutants, and is averaged per species rather than per mutated gene in this case. **(C)** Average caterpillar weight of all caterpillar species appearing in panels (A) and (B) excluding *H. zea,* divided by plant genotype on which they were fed. The weight of each caterpillar was normalized to its species WT values for that plant genotype. **(D-H)** Average caterpillar weight at the assays end. **(D)** *S. exigua*. **(E)** *S. eridania*. **(F)** *T. ni*. **(G)** *H. virescens*. **(H)** *M. sexta*. In panel (A), different letters indicate a significance of p<0.05 as determined in a Tukey HSD test. *H. zea* n= 45, *S. exigua* n=98, *S. eridania* n=107, *T. ni* n=*118, H. virescens* n=94, *M. sexta* n=54. In panel (B) different letters indicate significance of p<0.05 as determined in a Chi square test. *H. zea* n= 117, *S. exigua* n=135, *S. eridania* n=142, *T. ni* n=*129, H. virescens* n=135, *M, sexta* n=66. In panel (C) WT n=107, *cer1* n=77, *cer3* n=112, *kcs6* n=62, *far* n=113. In panel (D) WT n=20, *cer1* n=19, *cer3* n=17, *kcs6* n=20, *far* n=22. In panel (E) WT n=30, *cer1* n=16, *cer3* n=31, *far* n=30. In panel F: WT n=24, *cer1* n=19, *cer3* n=26, *kcs6* n=23, *far* n=26. In panel G: WT n=17, *cer1* n=16, *cer3* n=23, *kcs6* n=19, *far* n=19. In panel H: WT n=16, *cer1* n=6, *cer3* n=16, *far* n=16. WT = wildtype *N. glauca.*

When correlation of the normalized average caterpillar weight divided to the different plant genotypes was examined in relation to the wax components (Fig 5B-C), anabasine (Fig. 5A) and sugars (Fig. 5D), high correlation was found with the anabasine content (r^2^=0.765). This was in line with low caterpillar weight on *cer1* and *kcs6* leaves, possessing higher anabasine content than the other genotypes. An exception was the tobacco specialist *M. sexta*, which is expected to be less affected by anabasine (Huesing and Jones, 1987). To uncover the effects of the wax components regardless to the anabasine content, we correlated the mass of *M. sexta* caterpillars with the metabolites in a similar manner. Here, correlation with anabasine was poor (r^2^=0.007; Fig. 5E) whereas correlation with fatty alcohols rose drastically (R^2^=0.858; Fig. 5G), a possible explanation for caterpillars growing faster on the *far* fatty alcohol synthesis mutant. Caterpillars’ response when caged on individual leaves may differ from their response on whole plants, where the eaten leaf may be chosen based on parameters such as leaf age or water status. Therefore, to complement the experiment with six caterpillar species caged on individual leaves, we examined the effects of wax composition on the growth of *S. littoralis* on whole plants. Similar to the previous results, *S. littoralis* grown on *far* mutants were significantly heavier than those grown on all other lines, although caterpillars grown on *cer3* plants were similar to WT (Fig. 6A).

**Figure 5.**
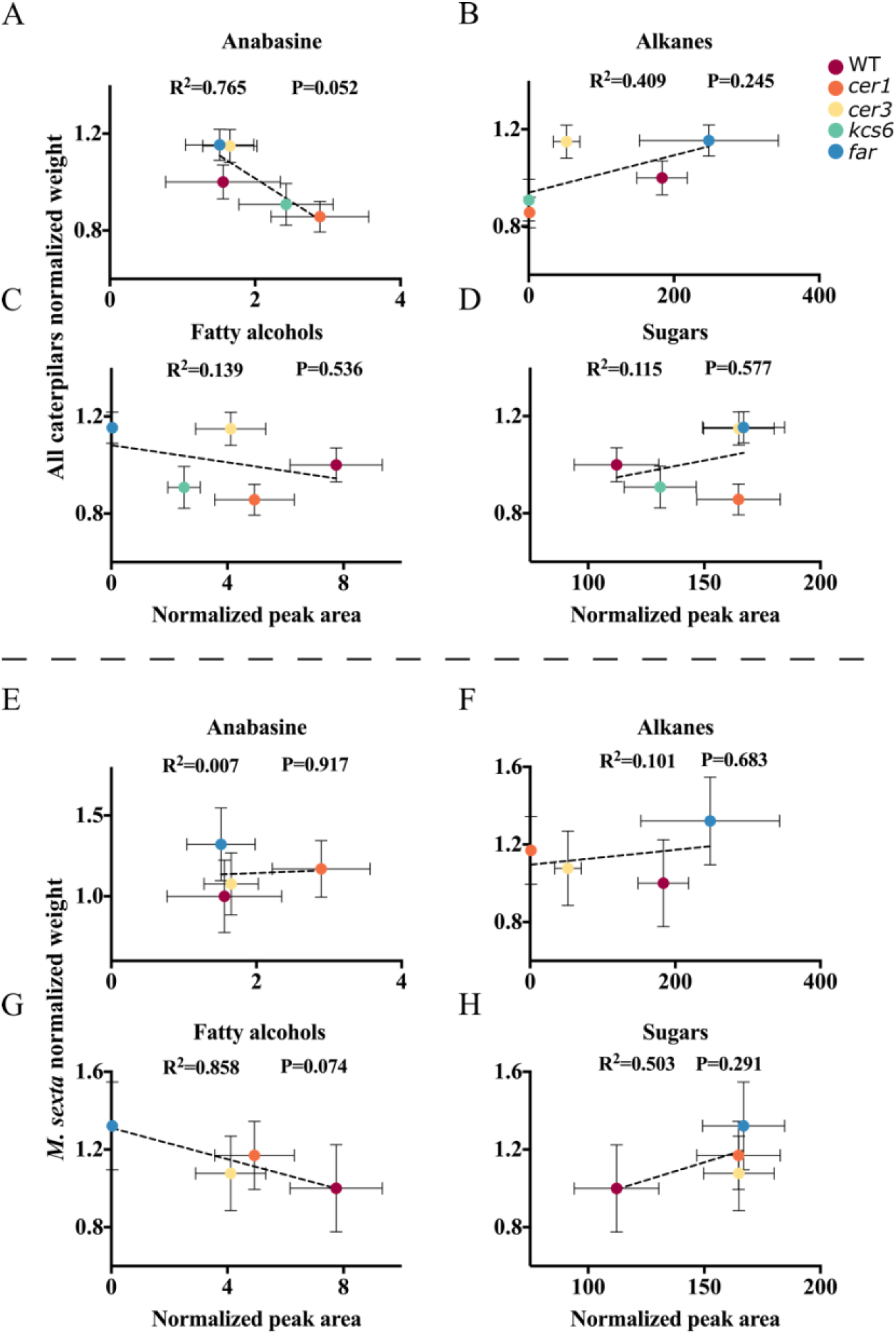
Correlation between wax and polar metabolite abundance and caterpillar weight. Each differently colored point indicates a genotype on which caterpillars were grown, and whose wax and polar metabolite composition were assessed (Fig. 3,S1). **(A-D)** Combined average weight and metabolite abundance of *S. exigua*, *S. eridania*, *T. ni*, *H. virescens*, and *M. sexta*. **(E-H)** Average weight and metabolite abundance of *M. sexta*. (A) and (E) correlation of anabasine content to caterpillar weight. (B) and (F) correlation of alkane content to caterpillar weight. (C) and (G) correlation of very long chain fatty alcohol content to caterpillar weight. (D) and (H) correlation of the average sum of five highly abundant sugars and caterpillar weight. The analyses in panels A-D were performed on WT and four wax mutants, whereas those in panels E-H did not include the *kcs6* mutant. Bars on the sides of points on the Y axis are standard errors of caterpillar weight whereas those on the X axis are standard errors of the metabolite abundance. R^2^ values as well as p values calculated by an F test for slope equality to 0 are presented in each panel.

**Figure 6.**
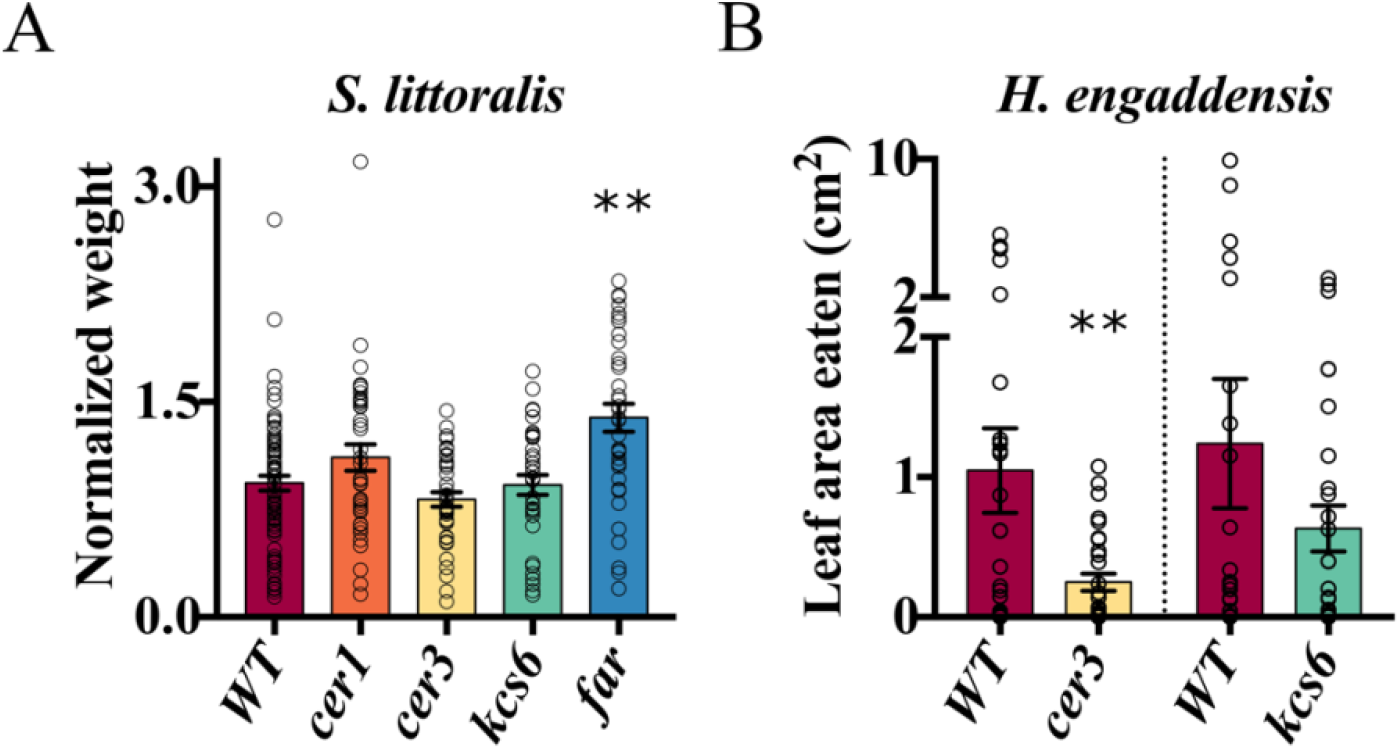
Effect of wax composition on chewing herbivores growth and feeding. **(A)** effects of wax composition on growth of *S. littoralis* caterpillars when placed on whole *N. glauca* plants. Caterpillar weight was normalized to each block’s average caterpillar weight. WT n=78, *cer1* n=38, *cer3* n=40, *kcs6* n=36, *far* n=37 Double asterisks indicate significance of P<0.01 as determined in a Dunnett’s test. **(B)** effects of wax composition on leaf area eaten by *H. engaddensis* snails when confined to a WT and a wax mutant leaf. Three independent replicates of 10 boxes with WT and *cer3* leaves and 10 boxes with WT and *kcs6* leaves were performed. Double asterisks indicate significance of P<0.01 as determined in a matched pairs t test. N=30. WT = wildtype *N. glauca.*

Next, we examined the effects of wax composition on *H. engaddensis* herbivory. These were chosen following observations of *N. glauca* growing in a natural setting, where snails consisted a major part of interacting herbivores. Unlike our original hypothesis, *cer3* leaves were eaten significantly less than WT leaves. The *kcs6* mutation did not affect snail herbivory (Fig. 6B).

### The influence of wax composition on phloem feeders

We next examined the impact on insect phloem feeding by performing *M. persicae* aphid reproduction assays. Aphids that were previously raised on pepper plants grew at significantly higher rates on *cer1* mutants, as seen at two time points as well as a significant interaction in a two-way ANOVA comparing the mutant line and day of measurement (Fig. 7A). Although we observed the same trend with aphids reared on tree tobacco, the result was not significant (Fig. 7B).

**Figure7.**
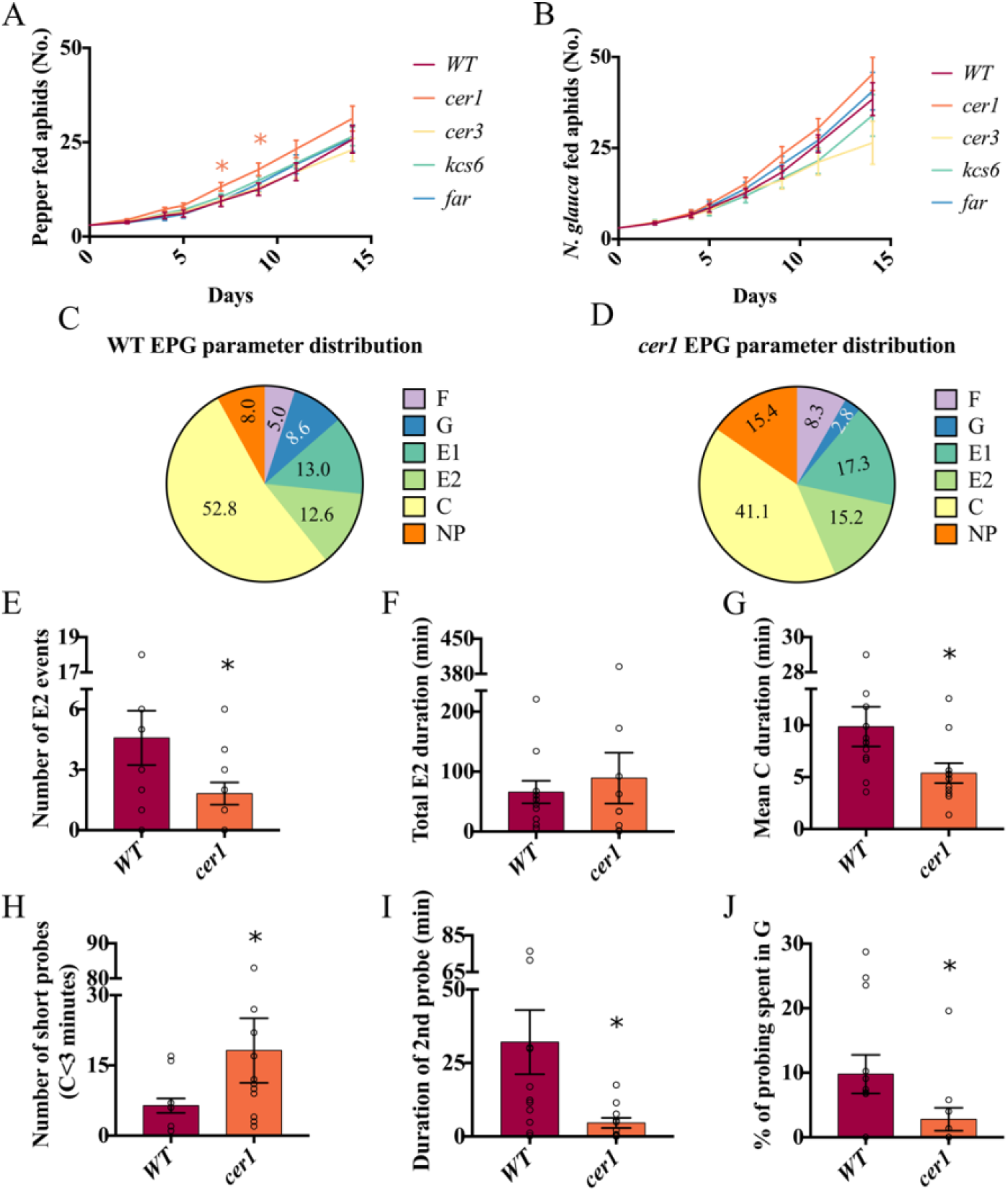
Effects of wax composition on *M. persicae* reproduction and activity. **(A-B)** Average number of aphids per cage, over time after transfer of three aphids from pepper (A) or *N. glauca* plants (B). Three separate replicates were performed with aphids from *N. glauca*, and five were performed with aphids from pepper. Aphid numbers in each cage were normalized to the replicate and day average, and then multiplied by the total average for that day. In panel (A), WT plants n=24, all other plants n=12. In panel (B), WT n=38, all other lines n=20. **(C-D)** Proportion of time spent by *M. persicae* performing different feeding activities on the leaves of WT **(C)** and *cer1* **(D)** plants. “NP” – non probing phase, when the aphids’ stylet is not in the leaf. “C” – activities during penetration which are mainly intercellular (probing). “E1” – salivation in a sieve element. “E2” – Ingestion of fluids from the phloem. “F” – derailed stylet mechanisms. “G” – xylem derived ingestion. **(E)** Total number of phloem ingestion events. **(F)** Total time spent in phloem ingestion (“E2”). **(G)** Mean duration of intercellular activity events (“C”). **(H)** Total number of probes, which lasted less than three minutes. **(I)** Average duration of the second probing event. **(J)** Proportion of the time spent in xylem ingestion out of the time spent probing. In panels (A-B), asterisks represent a significance of p<0.05 as determined by a Dunnett’s test. In panels (E-J), asterisks represent a similar significance in a Wilcoxon rank sum test. (C-J), WT n=12, *cer1* n=11. WT = wildtype *N. glauca*.

We next monitored the probing behavior of the nicotine-tolerant strain of *M. persicae* (Ramsey et al., 2007; Ramsey et al., 2014) on tobacco leaf surface using the electric penetration graph (EPG) technique. Employing EPG allowed us to better understand the underlying mechanism for the differences we observed, and assess whether wax composition affects aphid-feeding behavior in a way that could not be observed in reproduction assays. Data from the EPG is recorded as a pattern of electrical waves. These correlate to the different activities the aphid is performing, including non-probing (NP) when the stylet is not inserted to the leaf, probing (C) when the stylet is between cells searching for a fitting feeding location, sieve element salivation (E1), phloem ingestion (E2), xylem ingestion (G) and derailed stylet mechanisms (F). In addition to these, potential drops (PD) indicate stylet insertion to a cell after being in the intracellular region. Given that *cer1* was the only mutant on which aphids reproduced more in previous experiments, we compared aphid-feeding activity on WT and *cer1* plants. Unlike what may have been expected based on the reproduction assays (Fig. 7A), aphids on *cer1* leaves did not spend less time in the non-probing stage (Fig. 7C-D). Furthermore, although aphids on *cer1* leaves had significantly fewer phloem ingestion events (Fig. 7E), this was not reflected in the total time spent in the E2 phase (Fig. 7F). A parameter that was significantly shorter in *cer1* included the mean C duration (Fig. 7G), corresponding to time spent in the intercellular region, and a significantly higher number of short C events in *cer1* (Fig. 7H). The duration of the second probe was significantly lower in *cer1* (Fig. 7G) and the percentage of the probing time spent by the aphids in xylem ingestion, was lower in *cer1* mutants as well (Fig. 7J). The combination of these results points to aphids on *cer1* exploring the plant tissues with more ease.

### *B. tabaci* egg-laying on wax mutant leaf surface

The final interaction class that we examined involved *B. tabaci* egg-laying. Three experiments were performed in which we placed 20 *B. tabaci* in cages clasped onto leaves of WT and wax-mutant plants and counted the number of eggs laid 48 h later. Here, unlike in both chewing and phloem feeding herbivore experiments, where *far* and *cer1* mutants were more susceptible to herbivory. They showed a significantly lower number of eggs laid on them, whereas WT leaves were the most susceptible to egg laying. The alkaneless *kcs6* leaves were not significantly different as compared to WT ones (Fig. 8).

**Figure 8.**
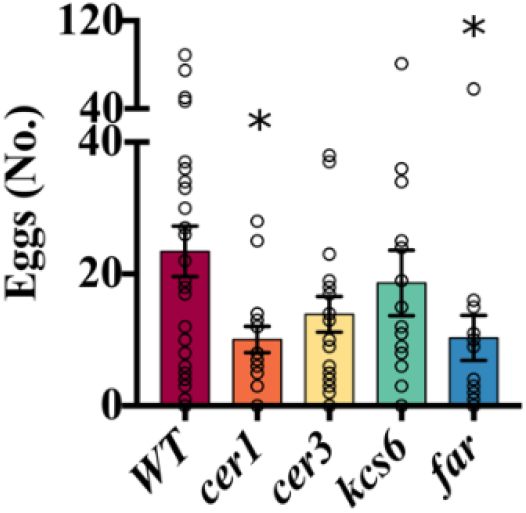
Effects of wax composition on the number of eggs laid by *B. tabaci*. Three separate replicates of the assay were performed. Asterisks indicate a significance of p<0.05 as determined in a Dunnett’s test. WT n=30, *cer1* n=16, *cer3* n=17, *kcs6* n=16 *far* n=16. WT = wildtype *N. glauca*.

## Discussion

Being the first barrier which the insect encounters, we expected epicuticular wax to have a strong effect on herbivory, whether it was to deter herbivores from feeding as seen in *Phyllotreta crucifera* and *Lipaphis erysimi* feeding on *Brassica Oleracea* or to induce eating, as has been shown in cases of *plutella xylostella*, *Pieris rapae* and *thrips tabaci* on *B. Oleracea* glossy mutants (Eigenbrode and Espelie, 1995). However, the effects of epicuticular wax on feeding of chewing herbivores was rather small when compared to those of secondary metabolites (see for example Gog et al., (2014). This is exemplified by *kcs6* mutants, which had values nearly identical to WT in the *H. engaddensis* and *S. littoralis* assays, and values slightly lower than WT in the multiple-species caterpillar assay. This wildtype-like insect growth was achieved despite *kcs6* mutant plants having almost no alkanes, as well as a reduced fatty alcohol content. In contrast, *far* mutants stood out in experiments with almost all caterpillar species, and especially in those where anabasine had less of an effect. *Far* mutants being more susceptible to leaf chewing is even more notable when considering that the *far* mutants have more wax than all other lines due to their elevated alkane levels. The negative correlation of *M. sexta* weight to fatty alcohol abundance also points to these fatty alcohols having an inhibitory effect on caterpillar growth. These results also may explain why *N. glauca* accumulates these alcohols since, when examining abiotic parameters in a previous study (Negin et al., 2021), we found that fatty alcohols *far* mutants have a slower rate of cuticular water loss, and are also more drought resistant on the whole plant level. Thus, this class of epicuticular waxes is likely to play a significant role in herbivory response rather than response to drought and other abiotic conditions.

In this study, we conducted insect bioassays using a population of *N. glauca* plants mutated in wax biosynthesis genes. An obstacle that we did not encounter in our previous research (Negin et al., 2021) with *N. glauca* was the different anabasine content of the mutant lines, which may have masked effects that may stem from the differential wax composition. Our *M. sexta* results point to the advantage of using specialist insects, which tend to tolerate the defensive metabolites of their host plants when examining effects of secondary metabolites, rather than using broad generalists that, although they manage to grow quite well in some cases, may hide the affects that we wish to investigate. Another approach we examined was the acclimation of the generalist *M. persicae* to feeding on *N. glauca* rather than pepper. In this case, we found that, although the growth rate of the aphids that were pre-acclimated on *N. glauca* was higher, the results were essentially similar (Fig. 7A, B).

The response of *M. persicae* was quite different compared to that of the chewing herbivores, with only the altered wax composition of *cer1* having a significant effect. It is interesting to note, that although *cer1* mutants have greatly reduced alkane content, there does not seem to be a correlation between aphid reproduction and alkane abundance. If alkane abundance were the underlying cause of higher aphid reproduction on *cer1* plants, we would expect to see similar results in the even more severe *kcs6* mutants as well as an intermediate phenotype in the greatly alkane reduced *cer3* mutants. This distinction is relevant because *cer1,* in addition to having an extremely altered epicuticular wax composition also has an altered cutin composition, with a significant increase in ω-hydroxylated fatty acids and a reduction in phenol content (Negin et al., 2021). In our previous study we found that, in cases where epicuticular wax strongly altered the plant’s response (such as in cuticular water loss or drought recovery assays), *kcs6* and *cer1* mutants displayed very similar results, whereas in cases where the effect was cutin-related (such as photosynthetic efficiency) *cer1’s* phenotype was altered but *kcs6’s* phenotype was not. For these reasons we therefore concluded that disruption of cutin composition increases aphid growth, whereas mutations affecting wax alone do not.

When investigating the *cer1* phenotype using EPG assays, we observed results that were similar to the growth assays. In case of epicuticular wax preventing aphids from inserting stylets to the leaf, we would expect both the time spent in the non-probing phase to be higher in WT and the time to the first probe to be longer. When examined, aphids on *cer1* leaves spent almost double the time in the non-probing phase (15.4% compared to 8%), although the time to the first probe was not significantly different. Additionally, although aphids on *cer1* plants had many more, and shorter C events, which are related with both epidermal and mesophyll factors, this did not prevent them from reaching a higher duration of phloem fluid ingestion (E2) in a significantly smaller number of E2 events. A larger number of shorter C events suggests that aphids on *cer1* leaves were choosier and probably explored the plant tissues with more ease. The trend of longer and less disturbed E2 events also may partly explain the higher aphid reproduction rates on *cer1* plants.

Non-feeding interactions of insects with the plant are affected not only by the surface’s chemical composition (Udayagiri and Mason, 1997) but by the leaf surface structure as well (Uematsu and Sakanoshita, 1989; Rahim Khan et al., 2011). In our oviposition assay, chemical composition did not correlate with the number of eggs laid, which could be better explained by the alterations in the surface structure. In this context, it is interesting to note that the intermediate number of eggs laid on *cer3* leaves may correlate with *cer3* wax crystal structure being similar to WT but more sparsely distributed (Negin et al., 2021). *Kcs6* plants stand out here, and their close to WT values may be related to the fact that, although their wax crystal structure is extremely altered, it maintains a rather organized distribution unlike the random one seen on *far*, *cer1* and *cer3* leaves (Negin et al., 2021).

Most if not all studies investigating effects of epicuticular wax on insect interaction focused on one or two altered wax compositions, one or two insects, and one interaction type [for example, see (Edwards, 1982; Städler and Buser, 1984; Daoust et al., 2010)]. This leads of course to very different and contradictory results. If indeed chewing insects are affected by chemical composition, phloem feeders by the ease to penetrate the cuticle, and non-feeding interactions affected in some cases by surface morphology, this may explain the differences between these assays.

An additional factor affecting plant-herbivore interaction is co-evolution. While abiotic factors are consistent, and a plant that developed a glaucous appearance to reduce UV damage will remain protected, this is not the case with herbivores. Almost every plant that has potent secondary metabolites protecting it also has an insect specialists that evolved to have a higher tolerance to these metabolites (Cornell and Hawkins, 2003; Ali and Agrawal, 2012). This may be seen in the tobacco specialist *M. sexta* (Self et al., 1964) used in this study, the resistance of the *Danaus plexippus* (monarch butterfly) to cardenolide toxins (Petschenka and Agrawal, 2015), the resistance of *Tequus sp.* (sawflies) to glycoalkaloids from the wild potato species *Solanum commersonii* (Altesor et al., 2014), and numerous other interactions. This co-evolution means that effects of metabolites on insect herbivory, even when the synthesis of these compounds evolved to protect the plant from herbivory, may vary widely between insect species. The use of several insect species as in this study is hence most valuable for investigating plant-herbivore interactions.

Our study here points to a significant contribution of epicuticular wax composition to plant-herbivore interaction. Yet, plant-insect interactions are highly intricate as they are greatly affected by co-evolution and are very species and interaction type specific. However, as a generalization we may say that epicuticular wax load and alkane abundance do not contribute to resistance against chewing herbivores, whereas very long chain fatty alcohols have an inhibitory effect on lepidopteran growth. In generalist species this effect is somewhat masked by the more drastic growth inhibition caused by anabasine in *N. glauca*. However, it is more apparent when the interaction is examined in *M. sexta*, a nicotine tolerant herbivore that was shown to be tolerant to anabasine as well (Huesing and Jones, 1987). *M. sexta* was the lepidopteran species that had the fastest growth and was least affected by anabasine in our assay, as seen in the correlation of anabasine to weight gain. In contrast to caterpillars, *M. persicae* are unaffected by fatty alcohol content. Instead, they reproduce at higher rates when the cutin composition is disrupted. Once again, there seems to be little effect of the total epicuticular wax load in this context. Non-feeding interactions examined in this study seemed to be dependent on leaf surface morphology, and less on the chemical composition of the leaf. In the case of *B. tabaci*, dense wax crystal coverage contributed to larger numbers of eggs being laid. Our findings suggest that the specific composition of the epicuticular wax was in part shaped by interactions with herbivores and still plays a contributory role in these interactions.

