## Supplemental figure 1 for "Fatty Alcohols, a Minor Component of the Tree Tobacco Surface Wax, Reduce Insect Herbivory"

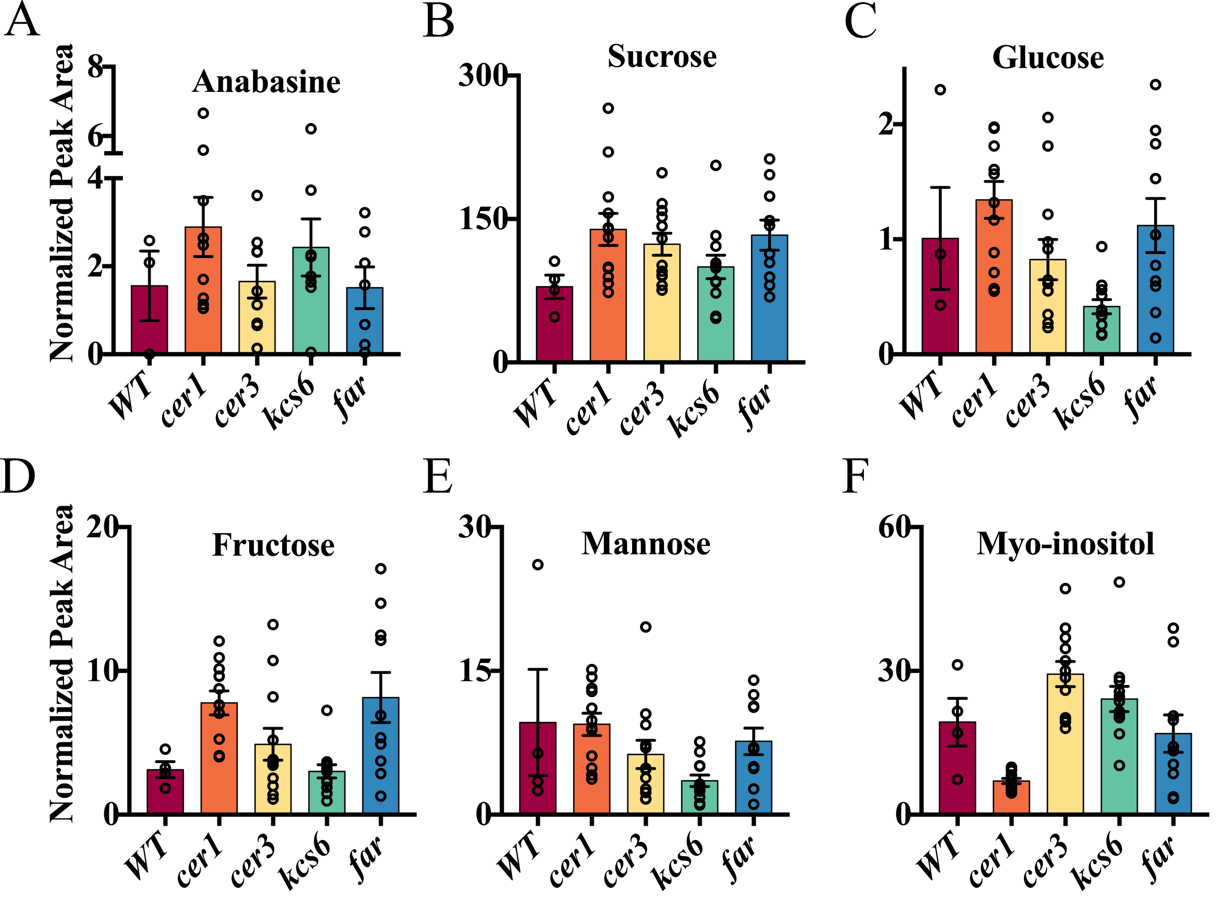

**Figure S1.** Relative abundance of prominent polar metabolite in leaves of WT and four epicuticular wax metabolism mutant lines of *N. glauca*. (**A)** Anabasine. (**B)** Sucrose. (**C)** Glucose. (**D)** Fructose. (**E)** Mannose. (**F)** Myo-inositol. Dunnett’s test was used to examine statistical significance. For anabasine, WT n=3, *cer1* n=9, *cer3* n=9, *kcs6*=8, *far* n=7. Sugars WT n=4, *cer1*, *cer3*, *kcs6* n=12, *far* n=10. Three independent mutant alleles were analyzed for each gene and then pooled for this analysis. WT = wildtype *N. glauca.*
